# Mutual information for detecting multi-class biomarkers when integrating multiple bulk or single-cell transcriptomic studies

**DOI:** 10.1101/2024.06.11.598484

**Authors:** Jian Zou, Zheqi Li, Neil Carleton, Steffi Oesterreich, Adrian V. Lee, George C. Tseng

## Abstract

**Motivation:** Biomarker detection plays a pivotal role in biomedical research. Integrating omics studies from multiple cohorts can enhance statistical power, accuracy and robustness of the detection results. However, existing methods for horizontally combining omics studies are mostly designed for two-class scenarios (e.g., cases versus controls) and are not directly applicable for studies with multi-class design (e.g., samples from multiple disease subtypes, treatments, tissues, or cell types).

**Results:** We propose a statistical framework, namely Mutual Information Concordance Analysis (MICA), to detect biomarkers with concordant multi-class expression pattern across multiple omics studies from an information theoretic perspective. Our approach first detects biomarkers with concordant multi-class patterns across partial or all of the omics studies using a global test by mutual information. A post hoc analysis is then performed for each detected biomarkers and identify studies with concordant pattern. Extensive simulations demonstrate improved accuracy and successful false discovery rate control of MICA compared to an existing MCC method. The method is then applied to two practical scenarios: four tissues of mouse metabolism-related transcriptomic studies, and three sources of estrogen treatment expression profiles. Detected biomarkers by MICA show intriguing biological insights and functional annotations. Additionally, we implemented MICA for single-cell RNA-Seq data for tumor progression biomarkers, highlighting critical roles of ribosomal function in the tumor microenvironment of triple-negative breast cancer and underscoring the potential of MICA for detecting novel therapeutic targets.

**Availability:** https://github.com/jianzou75/MICA

## 1. Introduction

Biomarker detection provides information for early disease diagnosis and is a critical element in biomedical research [Liu et al., 2020]. Integration of data from multiple cohorts is a common approach to improve reliability and statistical power of biomarker detection. If a biomarker demonstrate a similar pattern across multiple studies, it provides robustness and high likelihood of success in subsequent translation and clinical applications. In transcriptomic analysis, differential expression (DE) analysis stands as the predominant method for identifying biomarker expression pattern within individual studies [Costa-Silva et al., 2017, Conesa et al., 2016, McDermaid et al., 2019]. However, the majority of DE techniques are tailored for two-class scenarios (e.g., case versus control), faltering in multi-class scenarios. Popular methods such as limma [Ritchie et al., 2015], although capable of handling multiple classes, primarily offer statistical tests for aggregated differential information in a global sense rather than considering the expression patterns. This limitation highlights a paucity of methods adept at delineating multi-class expression patterns.

To address the integration of omics analysis results from multiple cohorts, two popular approaches emerge in the literature: combining p-values and combing effect sizes. The former has been widely discussed. For example, Fisher’s method sums up the log-transformed p-values, and each p-value is assumed to follow standard uniform distribution under the null hypothesis. In addition to Fisher’s method, Stouffer [Stouffer et al., 1949], minimum p-value [Tippett et al., 1931], higher criticism [Donoho and Jin, 2004], and adaptive Fisher method [Li and Tseng, 2011] have been developed under this category and are widely used in the omics study integration, such as GWAS [Begum et al., 2012], transcriptomics [Tseng et al., 2012], and methylation [Smith et al., 2018]. Random effects models [DerSimonian and Kacker, 2007], an example of the latter approach, decompose each study’s observed treatment effects into the actual effect size and the study-specific noise. These methods, however, have limitations to combine multi-class differential information. P-value combination methods focus on significance without considering multi-class patterns, while effect size combination is restricted to two-class scenarios. To our knowledge, the min-MCC method [Lu et al., 2010] is the only established approach for detecting concordant multi-class biomarkers across multiple studies, The method, however, has two major drawbacks on overlooking the situation when only partial studies share the multi-class pattern and not distinguishing between cases where all pairs of studies have a uniformly low concordance and cases where only one pair has a very low concordance.

To address these challenges, we introduced Mutual Information Concordance Analysis (MICA), a novel two-stage framework for multi-class biomarker detection combining multiple studies from the perspective of information theory. The first stage employs the generalized mutual information with one-sided correction (*gMI*_+_) to overcome the aforementioned drawbacks. The second stage involves a post-hoc pairwise analysis to identify studies sharing the concordant expression pattern. In 2024, where sequencing studies are ubiquitous, having a method like MICA can be a powerful tool for integrating datasets and enhancing the detection of robust biomarkers. We focus on bulk and single-cell transcriptomic applications in this paper but the method are readily applicable to other omics data types.

As a visual demonstration, Fig 1A shows three example genes *Amacr, Pole*4 and *Mcrip*2 detected by MICA to have concordant multi-class (WT: wild type mice; LCAD: LCAD mutated mice; VLCAD: VLCAD mutated mice) pattern across all or partial studies (tissues) (enclosed by red rectangles) while *Mrpl*51 is not detected due to heterogeneous patterns in all four tissues. Post-hoc pairwise analysis in the second stage then determines the studies (enclosed by yellow and blue triangles) that contribute to such concordance for the genes identified in the first step. Specifically, all four tissues share the same multi-class expression pattern in *Amacr*. Only brown fat, heart and liver tissues but not skeletal tissue share the same multi-class expression pattern in *Pole*4. Interestingly, in *Mcrip*2 gene, brown fat and liver share one concordant pattern, while Heart and Skeletal share a different concordant pattern.

**Figure 1.**
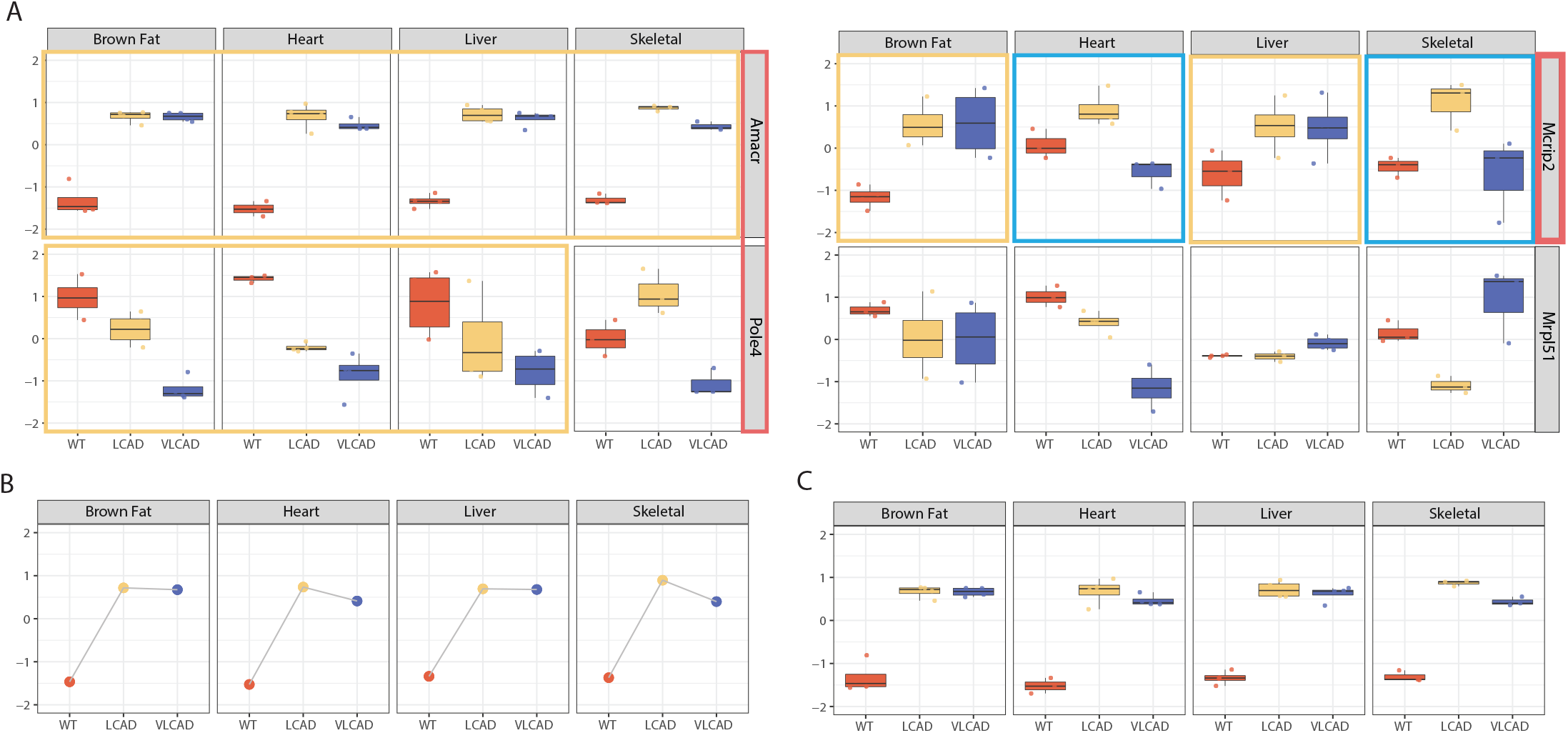
Depiction of the MICA Framework. The MICA framework was illustrated using the mouse metabolism dataset. (A) The application of the MICA framework to different gene types. Genes *Amacr, Pole*4, and *Mcrip*2 that exhibited consistent patterns across studies were initially identified by a global test using *gMI*+ (highlighted by the red triangle). Subsequent post-hoc tests using *MI*+ (highlighted by yellow and blue triangles) detected the studies sharing this consistency for each gene. (B) The scenario without replicates for each class within each study, displaying the median expression of the *Amacr* gene within each class and tissue. (C) The scenario with multiple replicates for each class within each study, displaying the expression of all samples for the *Amacr* gene.

The paper is structured as follows. In Section 2, we firstly review the existing method multi-class correlation (MCC) and min-MCC [Lu et al., 2010], followed by a reappraisal from an information theoretic perspective, where we demonstrate improved properties of the MICA framework. A simulation study and three real-world bulk and single-cell transcriptomic applications (Section 3) are conducted to compare min-MCC and MICA. Conclusions and discussions of MICA are included in Section 4.

## 2. Methods

We assume input data to contain *K* classess (*K* ≥ 2) for detecting multi-class patterns in *S* transcriptomic studies for integration. For simplicity, we skip subscript of genes and denote *x*_*ski*_ as the gene expression for one gene in study *s* (1 ≤ *s* ≤ *S*), class *k* (1 ≤ *k* ≤ *K*), and sample *i* (1 ≤ *i* ≤ *n*_*sk*_). For clarity, when discussing the two-study scenario (i.e., *S* = 2), we employ *x*_*ki*_ to represent the gene expression in study *X*, and similarly *y*_*ki*_ for study *Y*.

### 2.1 Multi-class correlation (MCC)

We start from the case of two studies (*S* = 2) with expression vectors *X* and *Y*. We first consider the simplest case wherein *n*_*sk*_ = 1 for all the studies *s* (1 ≤ *s* ≤ *S*) and the classes *k* (1 ≤ *k* ≤ *K*) (Fig 1B). Under this circumstance, the intuitive strategy for calculating the concordance (correlation) between study *X* and *Y* utilizing Pearson correlation is as follows:

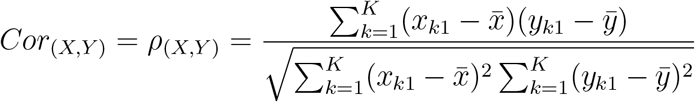

where 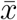 and 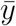 respectively denote the means of *x*_*k*1_ and *y*_*k*1_ for all 1 ≤ *k* ≤ *K*.

When there are replicates within each class from each study (*n*_*sk*_ > 1, Fig 1C), Pearson correlation is no longer viable. For study *X*, the observed gene expression *x*_*ki*_ is assumed to be obtained from 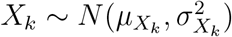, where *X*_*k*_ ╨ *X*_*k*_*′* (∀ *k* ≠ *k*^′^). Therefore, study *X* can be naturally defined as a mixture distribution of *X*_*k*_ (*k* = 1: *K*), where each class is assumed to be equally weighted.

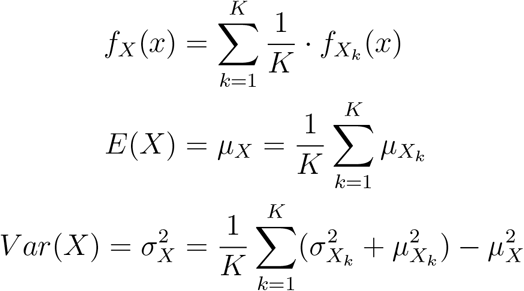

Study *Y* is similarly defined, and *Y*_*k*_ is independent with *X*_*k*_. The above-mentioned parameters can all be directly estimated from the data.

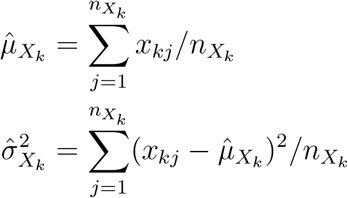

Multi-class correlation (MCC) is therefore defined as

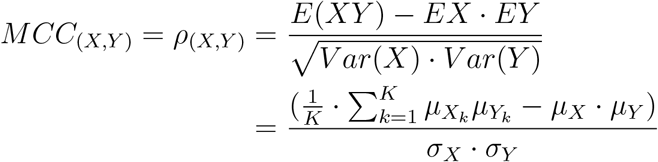

For multiple *S* studies (*S* > 2), min-MCC [Lu et al., 2010] is then defined a s t he m inimum value of MCC statistics across all the pair-wise study combinations:

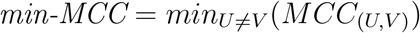

The hypothesis test *HS*_*A*_ for min-MCC to detect concordant expression pattern across all *S* studies is *H*_0_: ∃ *ρ* _*ij*_ ≤ 0 vs. *H*_*A*_: ∀ *ρ* _*ij*_ > 0, where *ρ* _*ij*_ represents the measurement of concordance in the multi-class pattern between study *i* and *j*. In addition to computational burden when *S* is large, min-MCC has two drawbacks. First, it neglects the situation when the concordant multi-class pattern only exists in partial studies due to its stringent requirement for consistency across all studies. Second, it cannot differentiate between scenarios where all study pairs have uniformly low concordance and scenarios where only one pair has very low concordance, which can lead to misinterpretations.

### 2.2 Mutual information concordance analysis (MICA)

To overcome the issues above, we revisit this problem from the aspect of information theory. We assumed *X* and *Y* to be jointly bivariate normal and denote *Z* and *Z* ^╨^ as the bivariate random variables when *X* and *Y* are correlated (*ρ* ≠ 0) or no correlation respectively.

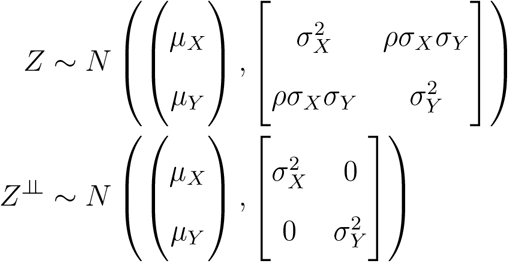

Therefore, we can define the mutual information between *Z* and *Z* ^╨^ as

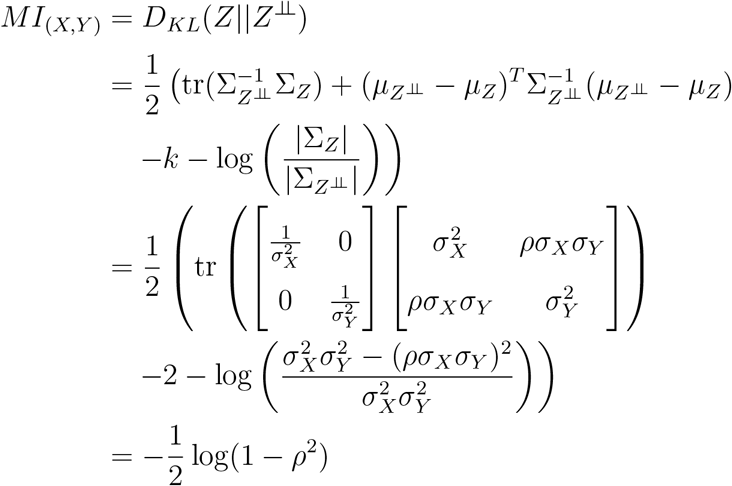

*D*_*KL*_ means the Kullback-Leibler divergence, and *ρ* is exactly the MCC between *X* and *Y*. To be consistent with MCC and limits to the positive correlation, we define the one-sided corrected mutual information (*MI*_+_) as

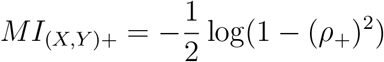

where *ρ* _+_ = *ρ* · 𝟙 _*ρ*>0_.

In the two-study scenario, we can find that *MI*_+_ is equivalent to MCC, but it is more straightforward to generalize to more than two studies. For *S* studies, we have *Z* ∃ *N* (***μ*, Σ**), *Z*^+^ ∃ *N* (***μ*, Σ**^+^), and *Z* ^╨^ ∃ *N* (***μ*, Σ** ^╨^), where

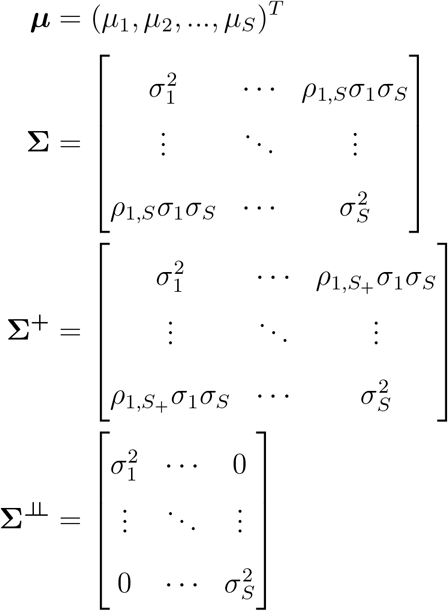

Therefore, we can define the concordance measurement for multiple studies, which is the generalized mutual information (*gMI*), also known as total correlation [Watanabe, 1960].

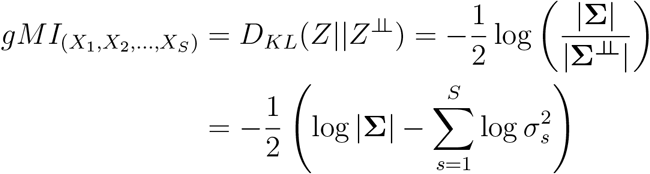

Similarly, to only consider the positive concordance, we define the generalized one-sided corrected mutual information (*gMI*_+_) as

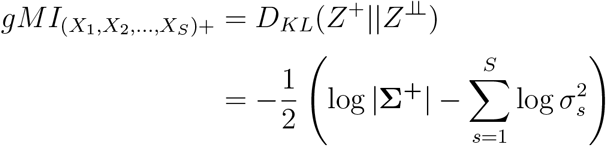

### 2.3 Procedure of concordant biomarker detection

Based on the generalized mutual information above, the mutual information concordance analysis (MICA) is developed in two steps.

#### 2.3.1 Global test for concordant biomarker detection

In the first step, we first deploy the generalized one-sided corrected mutual information (*gMI*_+_) to ascertain if a gene exhibits concordant multi-class pattern across multiple studies. This determination hinges on the hypothesis test, namely *HS*_*B*_, *H*0: ∀ *ρ* _*ij*_ ≤ 0 vs. *H*_*A*_: ∃ *ρ* _*ij*_ > 0. The permutation test (see Section 2.3.3) is employed for assessing *p*-values and *q*-values of this global test for each gene.

#### 2.3.2 Post-hoc test to detect subset of studies with concordant multi-class pattern

If the null hypothesis in the global test is rejected, we proceed to identify the subest of studies with concordant multi-class pattern. Specifically, we select the largest subset of studies where every pair of studies in it shows a significant p-value, indicating concordance. For this purpose, we employ the one-sided corrected mutual information (*MI*_+_) to examine all feasible pairs of studies (*i, j*). This analysis is conducted under the hypothesis setting *HS*_*C*_ for the study pair *i* and *j* by *H*_0_: *ρ* _*ij*_ ≤ 0 vs. *H*_*A*_: *ρ* _*ij*_ > 0, with p-values inferred by permutation test (see Section 2.3.3).

#### 2.3.3 Permutation test for the four statistics

Permutation test is designed to obtain the significance levels for *MI*_+_ and *gMI*_+_ since an analytical solution is not achievable. We use *θ* to denote them for using permutation test to evaluate *p*-values and *q*-values. To compare with existing methods, we use the same permutation analysis for MCC and min-MCC.

1. Compute statistics *θ* _*g*_ for gene *g*.
2. Permutate the group label *B* times and calculate the permutated statistics 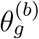, where 1 ≤ *b* ≤ *B*.
3. Calculate the p-value of *θ* _*g*_,

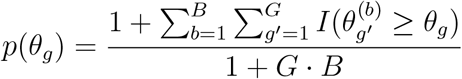
4. (If multiple genes are screened simultaneously) Obtain the p-values *p*(*θ* _*g*_) for each gene where 1 ≤ *g* ≤ *G*, and estimate q-values for gene *i* using Benjamini-Hochberg procedure. (*p* _(*j*)_ is ordered *j*-th p-value)

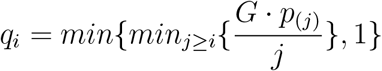

## 3. Results

In this section, we first applied MICA for simulations to evaluate the type I error and power of multi-class biomarker detection. The method is then applied to two bulk transcriptome applications: mouse metabolism-related studies [Lu et al., 2010], and estrogen treatment expression profiles [Li et al., 2023]. In the third application, we investigate the capability of MICA in single cell RNA-Seq data for tumor progression biomarkers detection. [Tokura et al., 2022, Wu et al., 2021a, Xu et al., 2021].

### 3.1 Simulation

We devised simulations involving five distinct types of genes from four studies (details in Supplement Table S1). Gene Type I represents perfect concordance with all four studies. In Gene Type II, studies 1, 2 and 3 show concordant expression. Gene Type III highlights pairwise concordance, showing agreement between studies 1 and 2 and a separate concordance between studies 3 and 4. Finally, Gene Type IV contains noises across all four studies, without any discernible pattern. There are 10 biological replicates within each class from each study, and the simulation is repeated for 500 times for evaluation. We then compare the performance of min-MCC and MICA in terms of type I error control and power.

MICA outperformed min-MCC in terms of signal detection power. For Gene Type I, where all studies were concordant, MICA achieved a detection rate of 0.836 against 0.638 for min-MCC at the p-value threshold of 0.05. In the more complex scenarios of Gene Types II and III, where only part of the studies were concordant, MICA maintained performance (0.748 in Gene Type II, 0.936 in Gene Type III), while min-MCC faltered (0.184 in Gene Type II, 0.174 in Gene Type III). Figure 2A-C provide a direct comparison of the respective powers of MICA and min-MCC at varying p-value thresholds across the three gene types. For the negative control, Gene Type IV, MICA exhibited an error rate of 0.058, slightly higher than the 0.048 error rate observed in min-MCC.

**Figure 2.**
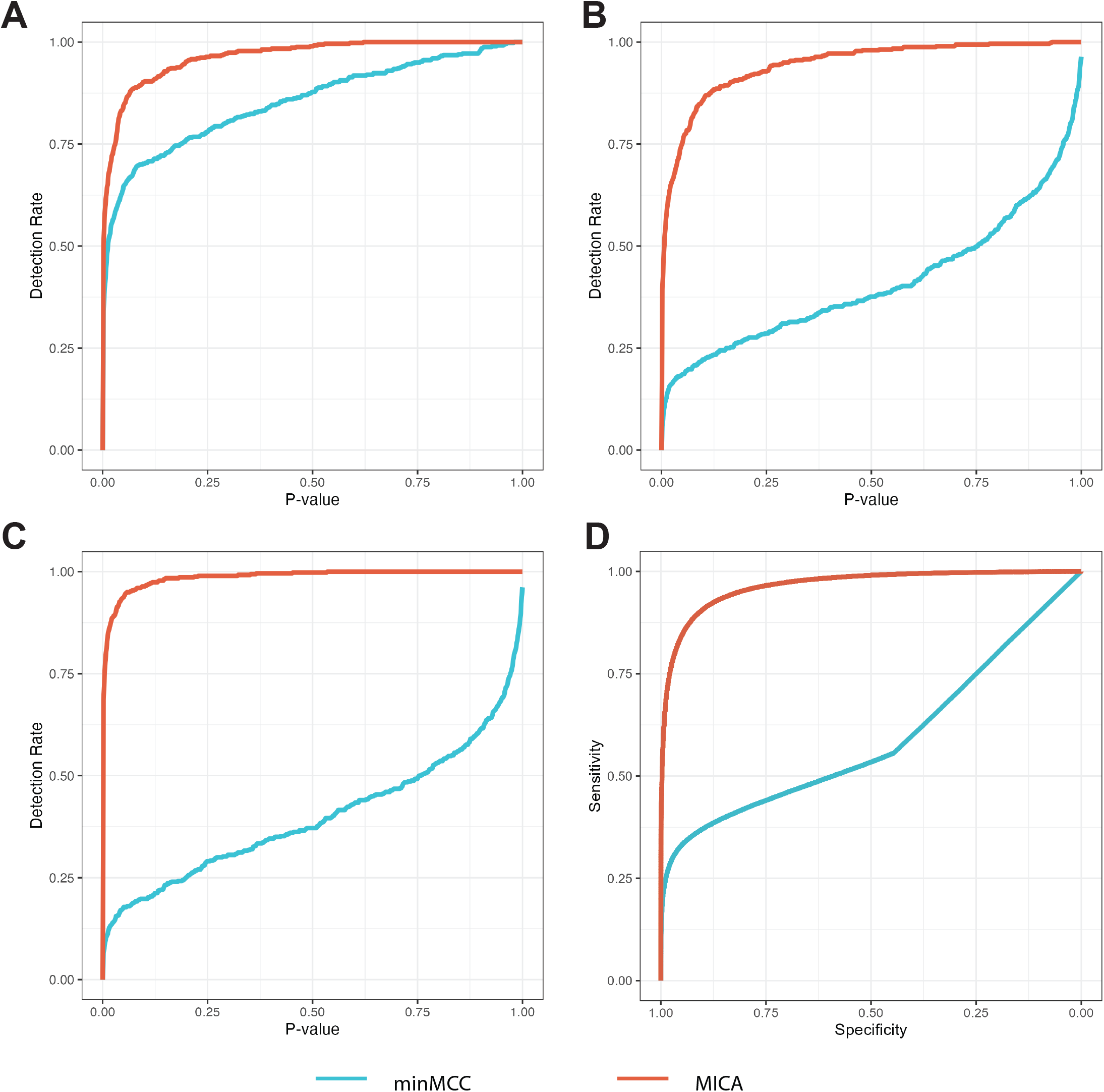
Comparative performance assessment of MICA and min-MCC via simulation. MICA consistently outperforms min-MCC in signal detection across Gene Types I-III. Genome-wide analysis further corroborates superior efficacy of MICA in biomarker identification. (A) Statistical power analysis in Gene Type I. (B) Statistical power analysis in Gene Type II. (C) Statistical power analysis in Gene Type III. (D) Aggregated ROC curves from 200 simulated datasets.

Following the assessment of individual genes, we expanded the simulation to encompass gene expression matrices for a genome-wide power comparison. We prepared 2000 genes expression for each dataset, distributed evenly across four gene types with 500 genes each. A total of 200 datasets was simulated for this analysis. After preparing the receiver operating characteristic (ROC) curves for each simulated dataset, the averaged area under the curve (AUC) of MICA was 0.97 (sd = 0.004), in contrast to 0.59 (sd = 0.02) for min-MCC. Figure 2D shows an ROC curve of the data aggregated across 200 simulated datasets, substantiating the superior performance of MICA. Employing a q-value threshold of 0.05, MICA achieved the sensitivity of 0.79 and the specificity of 0.97, whereas min-MCC has sensitivity and specificity at 0.23 and 0.99, respectively.

### 3.2 Application 1: mouse metabolism bulk transcriptomic studies

In this section, we applied MICA to the study analyzed in the min-MCC paper [Lu et al., 2010]. Bulk expression profiles are measured in mice with three genotypes (wild-type, LCAD knock-out, and VLCAD knock-out). LCAD deficiency is associated with impaired fatty acid oxidation, and VLCAD deficiency is associated with energy metabolism disorders in children. Microarray experiments were conducted on tissues from 12 mice (four mice per genotype) including brown fat, liver, heart, and skeletal. The expression changes across genotypes were studied, and genes with little information content were filtered out to have 4,288 genes remained for downstream analysis. Four samples were identified with quality defects and were excluded from further analysis.

A total of 730 concordant genes were identified through MICA analysis, while min-MCC only detected 245 concordant genes (q-value < 0.01), suggesting tissue heterogeneity. To evaluate the necessity of MICA, we classified the detected genes into three subsets: genes identified by min-MCC only (V), genes detected by min-MCC and MICA simultaneously (M1), and genes identified only by MICA (M2-M11). In the third subset, we classified genes into 10 modules based on post-hoc MICA results and clustered genes within the same module using *K*-means. The number of clusters was determined using the NbClust R package [Charrad et al., 2014].

Figure 3 and Supplement Figure S1 display the expression patterns for each gene module. Genes in Module V exhibited ambiguous expression patterns. Meanwhile, genes in Module M1, which were partitioned into two clusters, exhibited high concordance across all four tissues. We performed a QIAGEN Ingenuity Pathway Analysis (IPA) [Krämer et al., 2014] on genes in M1. Apart from the pathways known to be associated with LCAD and VLCAD [Nsiah-Sefaa and McKenzie, 2016], such as oxidative phosphorylation and acyl-CoA hydrolysis, metabolism and mitochondria-related pathways like arsenate detoxification, tetrapyrrole synthesis, and heme biosynthesis were also detected (Supplement Table S2). Additionally, genes in M1 showed more similar expression patterns in wild-type and VLCAD knock-out mice compared to LCAD knock-out mice, supporting previous findings that LCAD knock-out mice exhibit a more severe phenotype than VLCAD knock-out mice [Maher et al., 2010].

**Figure 3.**
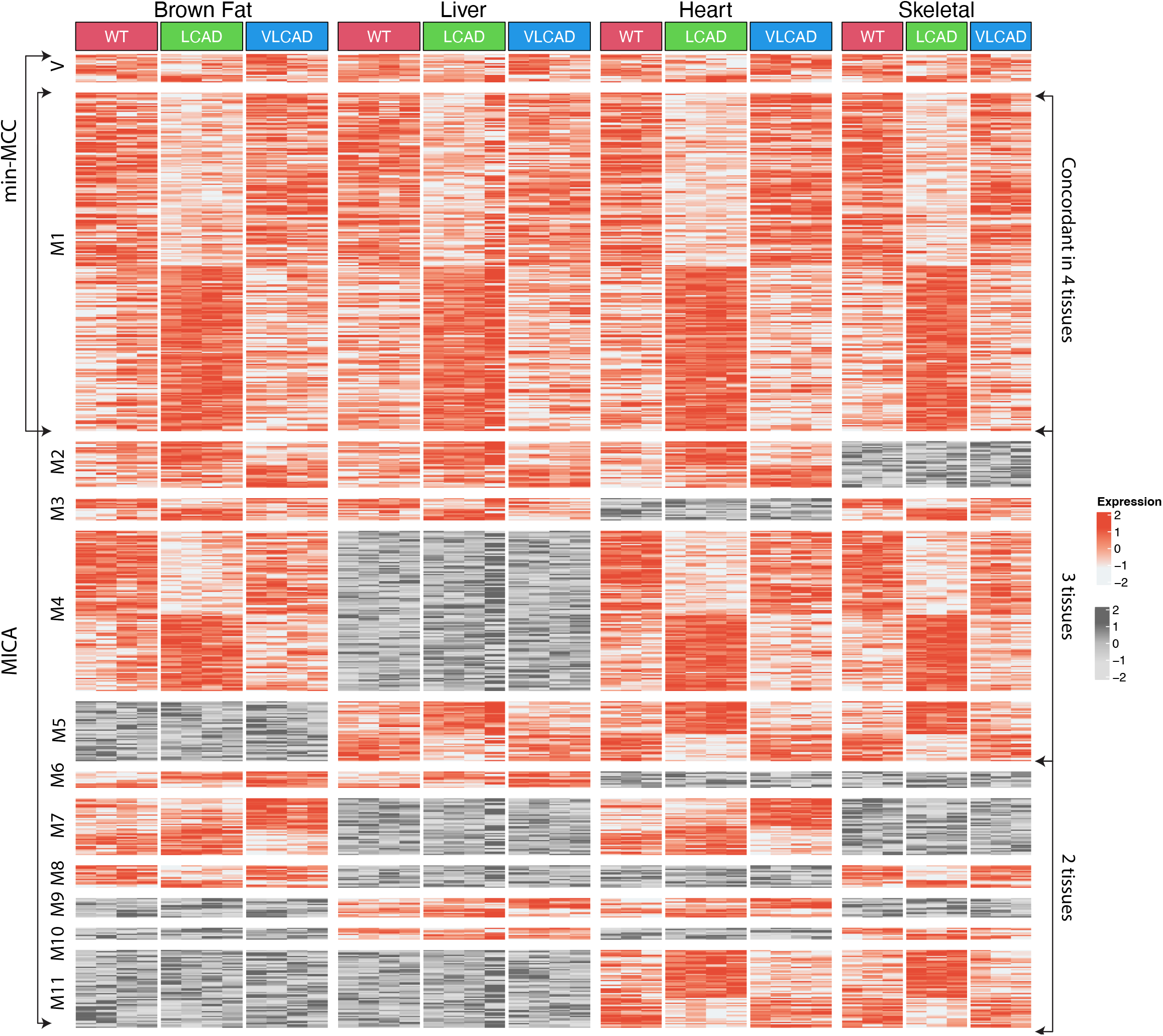
The heatmap of the gene expression patterns of different gene modules across four tissues in mouse metabolism data analysis. The rows represent the genes, and the columns represent the samples. V includes genes detected by min-MCC only, while M1 includes genes detected by min-MCC and MICA at the same time. The genes in M2-M11 were identified by MICA alone and categorized by the contributing studies using MICA post-hoc analysis. Studies that contribut-ed to the concordance are shown in red panel, while those that did not are shown in gray.

Modules M2-M11 demonstrated concordance pattern in a subset of tissues that were not detected by the min-MCC method. Among modules concordant in three tissues (M2-M5), Module M4 contained the largest number of genes (107 genes), showing impacts of LCAD and VLCAD knockouts in all tissues except for liver. *Blvrb* in Module M4 displayed the highest MICA statistic (MICA = 2.31, p-value = 0), although it was not identified by the min-MCC method (min-MCC = -0.71, p-value = 1) (Supplement Figure S2). *Blvrb* demonstrated lower expression in LCAD knockout samples in brown fat, heart, and skeletal tissues, but its expression was higher in the liver. Though *Blvrb* has no reported direct relation with LCAD and VLCAD, it is involved in metabolism, converting biliverdin to bilirubin in the liver [Consortium et al., 2017]. According to the Human Protein Atlas (proteinatlas.org) and the GTEx database [Lonsdale et al., 2013, Uhlén et al., 2015], *Blvrb* showed the highest gene expression in the liver among multiple tissues, indicating liver-specific functions not seen in the other three tissues.

The IPA analysis on genes in M4 (Supplement Table S2) emphasizes the distinct role of liver and the necessity to identify concordant pattern genes in a subset of tissues/studies. Specifically, 9 of the top 15 pathways, such as superpathway of methionine degradation and guanosine nucleotides degradation III, identified are related to metabolism, highlighting the role of liver.

In summary, MICA significantly outperforms min-MCC by identifying more concordant genes and uncovering tissue-specific gene expression patterns that min-MCC misses. This underscores the necessity of MICA for capturing the complexity of the partially concordant gene expression.

### 3.3 Application 2: bulk transcriptomic data in the EstroGene project

The EstroGene project [Li et al., 2023] focuses on improving the understanding of the estrogen receptor and its role in the development of breast cancer. It aims to document and integrate the publicly available estrogen-related datasets, including RNA-Seq, microarray, ChIP-Seq, ATAC-Seq, DNase-Seq, ChIA-PET, Hi-C, GRO-Seq and others, to establish a comprehensive database that allows for customized data search and visualization. Specifically, in this case, MICA can help identify genes that are consistently regulated by estradiol (E2) over different time points across multiple studies, which is critical for understanding the dynamics of estrogen receptor signaling in breast cancer.

In this subsection, we only considered studies that included gene expression data (microarray and RNA-Seq) and limited our analysis to the samples with estrogen receptor positive (ER+) treated with estradiol (E2) doses greater than 1nM for varying duration. We first combined the samples by cell line and sequencing technology. To further analyze the data, we then classified the treatment duration into three categories: short (< 6 hours), medium (≥ 6 hours and ≤ 24 hours), and long (> 24 hours). Finally, we normalized the data for the newly pooled studies using trimmed mean of M values (TMM) [Bullard et al., 2010] followed by ComBat [Johnson et al., 2007] with the study indication as a batch covariate. These steps resulted in three pooled studies: MCF7 microarray (25 samples in short treatment, 34 in medium treatment, and 7 in long treatment), MCF7 RNA-Seq (49 in short treatment, 62 in medium treatment, and 10 in long treatment), and T47D RNA-Seq (3 in short treatment, 22 in medium treatment, and 11 in long treatment). 1,983 genes were intersected across multiple platforms for downstream analysis.

We first validated the two well-established benchmark genes, *GREB*1 and *IL*1*R*1, which have been widely reported as E2 activated and repressed genes [Cheng et al., 2018, Rae et al., 2005, Schaefer et al., 2005, Lavigne et al., 2008]. Figure 4 revealed the up- and down-regulation of *GREB*1 and *IL*1*R*1 in MCF7 microarray and RNA-Seq studies. However, these trends were not observed in the T47D RNA-Seq study. Specifically, while T47D cells exhibited a decreasing trend in *IL*1*R*1 gene regulation from short to the combined medium and long durations (p < 0.05 from t-test), the trend reversed, showing an increase between medium and long durations (p < 0.05 from t-test), and no trend was observed in *GREB*1 gene (p = 0.85 from ANOVA test). This inconsistency is likely due to the inherent heterogeneity of breast cancer. Despite the inconsistency across all three studies, MICA evaluated the partial trend as concordant. As a result, MICA identified both genes as concordant with q-values of 0.01 and 0, while the min-MCC detected them with larger q-values of 0.03 and 0.06, respectively.

**Figure 4.**
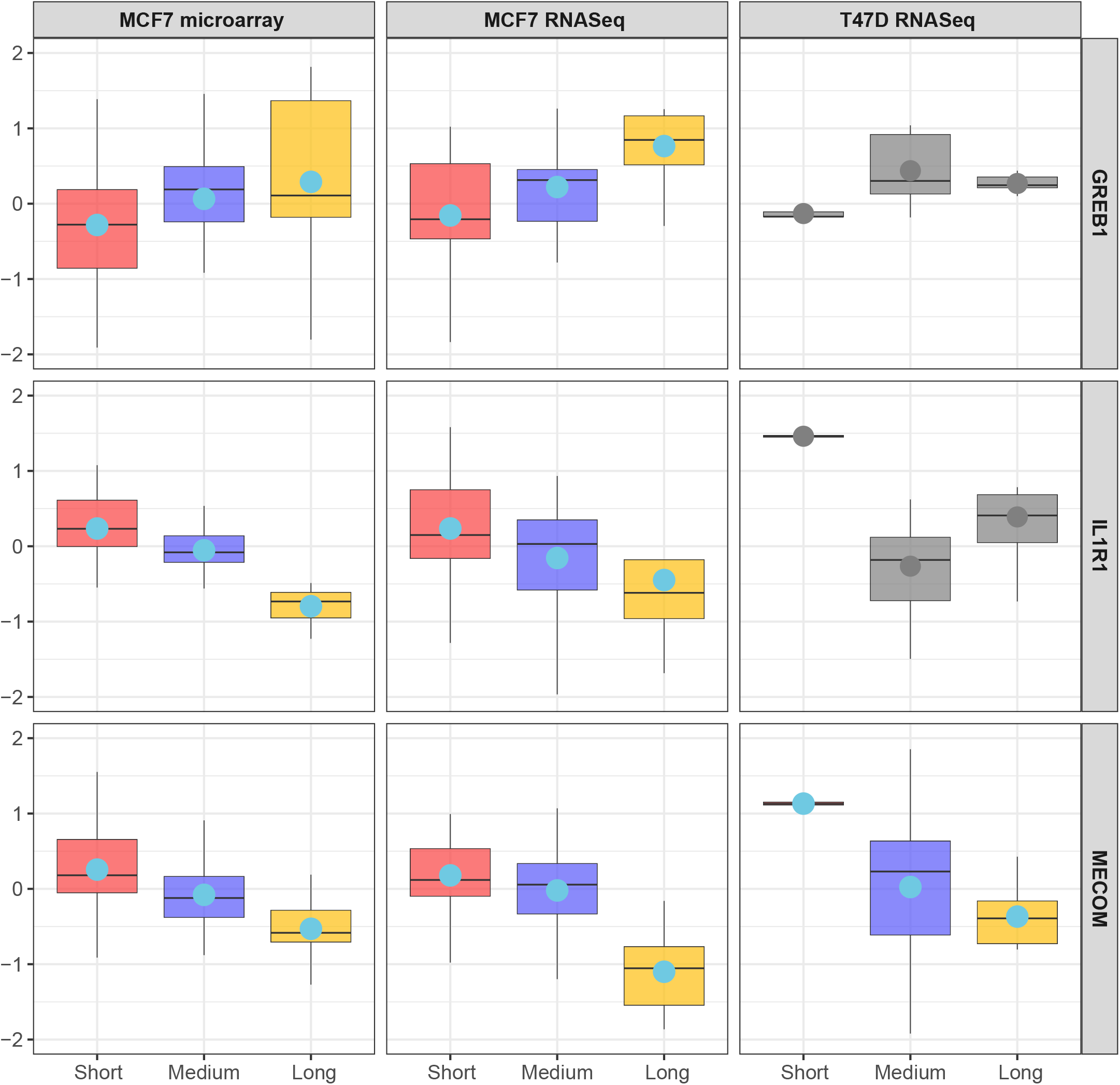
The expression patterns of *GREB1, IL1R1*, and *MECOM* across three data sources. *GREB*1 and *IL*1*R*1 are widely reported as E2 activated and repressed genes and were detected by MICA while failed to be identified by min-MCC. *MECOM* was the only gene detected by MICA and min-MCC simultaneously. The averaged expression is shown as a blue circle.

In addition to validating known markers, we are also able to detect novel biomarkers. For example, *MECOM* was the only gene identified by MICA and min-MCC with q-values = 0 simultaneously (Figure 4). Prior to our study, *MECOM* was not recognized as a biomarker for E2 treatment although it is known as a transcriptional regulator and oncogene. Indeed, when we analyzed 1,459 ER+ breast cancer patients in the Molecular Taxonomy of Breast Cancer International Consortium (METABRIC) database [Curtis et al., 2012], we observed that higher *MECOM* gene expression was associated with worse hazard ratio (HR) in terms of overall survival (HR = 2.27, p-value = 0.048) and relapse-free survival (HR = 3.34, p-value = 0.015).

To determine if this association is specific to HR+ tumors, we also performed survival analysis in other subtypes, including triple-negative breast cancer (TNBC) and HER2+ cohorts. In the TNBC cohort (n = 299), we did not observe a significant association (p-value = 0.18 for OS and p-value = 0.49 for RFS). Similarly, in the HER2+ cohort (n = 236), there was no significant association (p-value = 0.44 for OS and p-value = 0.90 for RFS). These findings suggest that the association of *MECOM* with survival outcomes is specific to HR+ tumors, which could strengthen the link between *MECOM* and endocrine response.

The potential mechanism of the clinical prognosis could partially be explained by the regulation of estrogen receptor, as we observed several consistent ER binding sites at transcription start sites (TSS) proximity from ChIP-seq data in the EstroGene website. The mechanistic link of *MECOM* to estrogen receptor and E2 treatment, however, needs further investigation.

In total, MICA identified 403 concordant genes (q-value < 0.05). To gain a deeper understanding of the upstream transcription factors associated with these genes, we applied LISA, an algorithm that uses chromatin profile and H3K27ac ChIP-seq data to determine the transcription factors (TF) and chromatin regulators related to a given gene set [Qin et al., 2020]. Among the top-ranked TFs (Supplement Table S3), *ESR*1 and *FOXA*1 are the TFs that have previously been reported to be associated with E2 [Chaudhary et al., 2017, Theodorou et al., 2013]. In addition, *SMC*1*A* and *CTCF*, the first two candidates, suggests a potential role of topologically associating domain (TAD) in the regulation of these gene [Rinzema et al., 2022, DeMare et al., 2013]. These findings revealed that the E2 response may involve gene regulation through chromatin looping mechanisms. Further experimental studies are needed to fully elucidate the underlying mechanisms.

### 3.4 Application 3: tumor progression biomarker detection in scRNA-seq breas cancer studies

In this subsection, we apply MICA to a scRNA-Seq dataset to compare three stages (*K* = 3) of triplenegative breast cancer (TNBC) progression using treatment-naive tissues: ductal carcinoma in situ (DCIS) (*N* = 5), primary tumor (*N* = 5), and lymph node metastasis (*N* = 2). Understanding the progression from DCIS, a precursor of invasive breast cancer, to primary tumors and eventually to metastatic disease is crucial for identifying biomarkers of tumor progression. The application of MICA in this case provides valuable insights into the molecular changes driving cancer metastasis, which is essential for developing targeted therapies and improving patient outcomes.

Data were obtained from three publications [Tokura et al., 2022, Wu et al., 2021a, Xu et al., 2021]. We implemented the scATOMIC [Nofech-Mozes et al., 2023] to annotate single cells to five cell types (B cell, CD4 T cell, CD8 T cell, macrophage and tumor cells) for downstream analysis. The distribution of cell types can be found in Supplement Table S4. Within each study, total count normalization was applied [Hao et al., 2023]. We treat the five cell types as independent studies (*S* = 5) and apply MICA to 6,644 genes after preprocessing. 2,703 genes exhibited concordant expression patterns across two or more cell types (q-value < 0.001). Notably, of the 86 genes associated with ribosomal functions, 82 exhibited concordance, which underscores the substantial role of protein synthesis in tumor progression.

In a further analysis, we aimed to identify the immune-tumor discordant genes, which exhibit concordant expression patterns across the first four tumor microenvironment cell types yet discordant patterns in tumor cells, as they progress from ductal carcinoma *in situ* (DCIS) to primary and subsequently to metastatic stages. To achieve this goal, we select from the 2,703 genes using criteria of any post-hoc pairwise p-values among immune cell types being less than 0.001 (i.e., all four immune cell types have concordant pattern to each other), and all pairwise p-values between a immune cell type and tumor exceeding 0.5 (i.e. all four immune cell types have discordant pattern to tumor). This analysis detected 198 genes (Supplement Table S5). Figure 5 illustrates the expression patterns of *RPS*15*A* and *RPS*25, the first two genes with the highest *gMI*_+_ statistics. The numbers of cells for each tumor type and each cell type are shown in the x-axis labels. Both genes are related to ribosomal functions, suggesting a hypothesis that protein synthesis is downregulated in immune cells as the tumor progresses, while it is upregulated in tumor cells.

**Figure 5.**
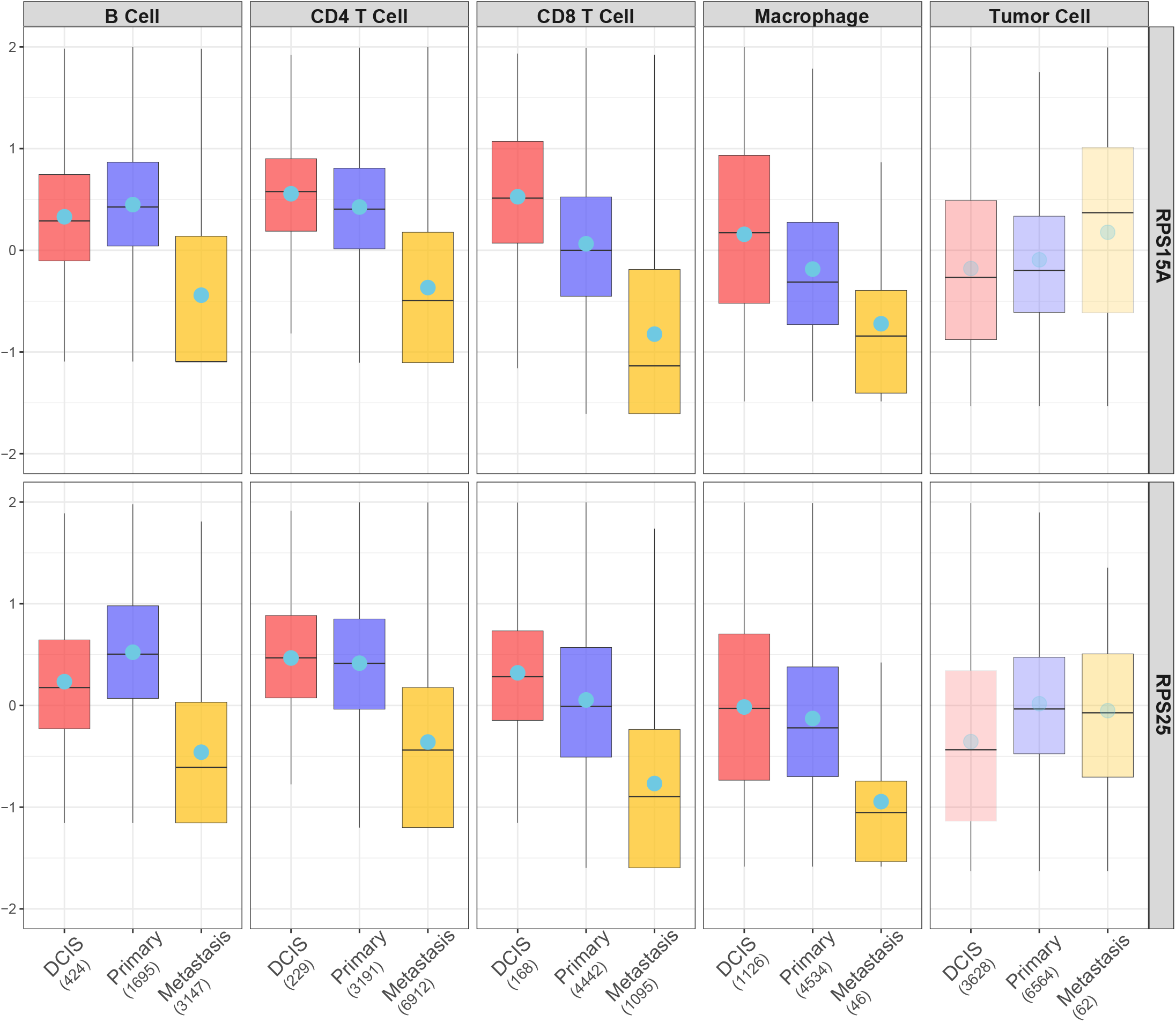
The expression pattern of *RPS*15*A* **and** *RPS*25. *RPS*15*A* and *RPS*25 are the top 2 immune-tumor discordant genes with largest *gMI*+ statistics. In immune cells, a decreasing expression trend is observed as the tumor progresses, whereas an opposing pattern is evident in tumor cells. The averaged expression is shown as a blue circle. The numbers of cells for each tumor type and each cell type are shown in the parentheses of x-axis labels

Gene set enrichment analysis [Wu et al., 2021b] of these immune-tumor discordant genes, conducted using the Gene Ontology (GO) knowledge base [Aleksander et al., 2023], revealed significant enrichment in two functional groups: cell junction and ribosome-related pathways (Figure 6). The former is closely associated with the tumor progression, such as altered cell adhesion and migration, while the latter confirms our earlier findings and underscores the importance of protein synthesis in the tumor microenvironment during the progression.

**Figure 6.**
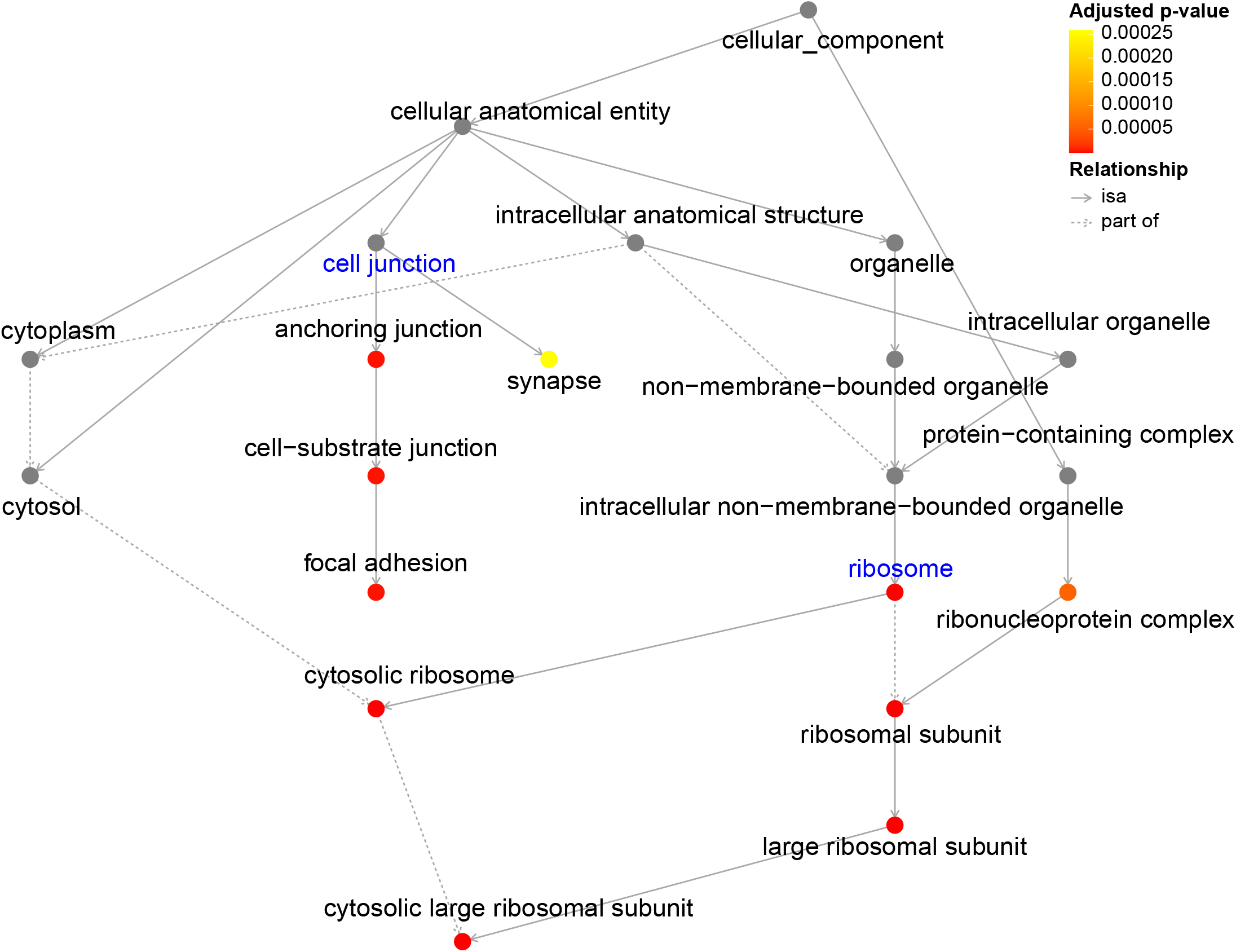
GO directed acyclic graph for enrichment analysis of immune-tumor discordant genes. Solid arrows (“isa”) denote that the term at the arrowhead is a subtype of the term at the tail, establishing a hierarchical relationship. Dashed arrows (“part of”) indicate that the term at the arrowhead is an essential part of the term at the tail, signifying structural inclusion. Dot colors represent the adjusted p-values from the enrichment analysis. Notably, cell junction and ribosome-related pathways are significantly enriched among the immune-tumor discordant genes.

## 4. Discussion and conclusions

Horizontal integration of multiple transcriptomic studies to identify disease biomarkers is an effective tool for accurate and reproducible detection [Cohn and Becker, 2003, Trikalinos et al., 2008]. To date, min-MCC is the only available method to detect the biomarkers with concordant multi-study multi-class expression patterns [Lu et al., 2010]. However, since min-MCC cannot identify the partially concordant biomarkers and is insensitive to the pairwise high concordance, we revisited this problem from the aspect of information theory and proposed a two-step framework MICA (Mutual Information Concordance Analysis). Both the simulation and real application results demonstrate the superiority of the MICA framework in selecting more informative biomarkers and elucidating underlying disease mechanisms towards translational research.

The three real applications contain a variety of biological and clinical scenarios and demonstrate wide applicability of MICA. In the mouse metabolism example, the biological objective is to detect biomarkers changed in wild type, LCAD or VLCAD mutation (*K* = 3) across four tissues (*S* = 4). Since biomarker pattern may different across different tissues, categorization of detected biomarkers in the heatmap of Figure 3 allows structured biological investigation. In the second EstroGene Project example, we investigate biomarkers with differential expression changes in short, medium and long treatment duration (*K* = 3) in three cell line-platform studies (*S* = 3). Ideally we expect similar expression pattern across studies but we indeed observe different multi-class pattern in different cell lines (e.g., GREB1 and IL1R1 in Figure 4). The third TNBC scRNA-seq example demonstrates an intriguing finding of tumor progression marker detection (DCIS, primary and metastatic tumor; *K* = 3) in five cell types of single cells (*S* = 5). The result identifies a set of biomarkers with concordant tumor progression pattern in the four immune-related cell types (i.e., B cell, CD4 T cell, CD8 T cell and Macrophage) while almost opposite pattern in tumor cells. We believe the wide range of applications not only demonstrate wide applicability of MICA but also will inspire its novel applications by other researchers.

One advantage of MICA is its scalable computing when *K, S* and biological replicate sample sizes increase. In the simulation of *K* = 3, *S* = 4 and 30 samples, the computing time for 2000 genes and 500 permutations takes 18.9 minutes using the high performance computing (HPC) with 50 threads parallel design. The current method performs analysis for each gene independently although the permutation scheme keeps gene dependence structure by permuting class labels when generating the null distribution for p-value assessment. An R package, namely MICA, and all programming code are available on GitHub for reproducing figures and results in this paper.

